# Controlling Strawberry Mites: Comparison of New Bio-Pesticides and Conventional Chemical Pesticides

**DOI:** 10.1101/677138

**Authors:** Jiahong Zhu, Zhuo Li, Zongjie Ren, Aocheng Cao, Dongdong Yan, Qiuxia Wang, Canbin Ouyang, Yuan Li

**Affiliations:** Department of Pesticides, Key Laboratory of Pesticide Chemistry and Application, State Key Laboratory for Biology of Plant Diseases and Insect Pests, Institute of Plant Protection, Chinese Academy of Agricultural Sciences, Beijing, 100193, China

**Keywords:** Bio-pesticide, Chemical pesticide, Toxicity test, Greenhouse test, Mechanism of pesticide action

## Abstract

**BACKGROUND:** *Tetranychus cinnabarinus* is one of the pest insects most severely influencing strawberry production. It has a high attack rate and causes severe economic losses. Laboratory toxicity tests and greenhouse experiments were carried out using 13 acaricides to determine their efficacy and potential mechanisms of action.

**RESULTS:** Abamectin showed the highest efficacy against *T. cinnabarinus* female mites; its LC_50_ value was 0.18mg L^-1^. Pyridaben, cyhalothrin, veratrine, and carbosulfan showed reasonably high efficacy; their LC_50_ values were 2.69 mg L^-1^, 3.94 mg L^-1^, 5.98 mg L^-1^, and 6.75 mg L^-1^, respectively. Less effective were hexythiazox and bifenthrin, their LC_50_ values were 9.82 mg L^-1^ and 19.09 mg L^-1^, respectively. Other acaricides such as spirodiclofen, chlorantraniliprole, chlorfenapyr, spinosad, and bifenazate did not show good efficacy. The status of female mites treated with avermectin, pyridaben, kanghebio and cyhalothrin changed significantly under a fluorescence microscope. There were no significant differences among female mites treated with spirotetramat, chlorantraniliprole, spinosad and bifenazate. Enzyme activity tests showed that Kanghebio and cyhalothrin obviously inhibited Ca^2+^-adenosine triphosphatase (Ca^2+^-ATPase), while veratrine and kanghebi obviously inhibited acetylcholinesterase (TChE) and monoamine oxidase (MAO). Cyhalothrinexerted an auxo-action on MAO. Greenhouse experiment indicated that abamectin showed the best efficacy, as well as the longest duration of efficacy, pyridaben, cyhalothrin, veratrine, and kanghebio followed, while carbosulfan, hexythiazox, and bifenthrin performed the worst.

**CONCLUSIONS:** Our study provided a scientific basis for chemical pesticides to be replaced by these and potentially other new bio-pesticides.

## 1. INTRODUCTION

*Tetranychus cinnabarinus* is one of the most important pest species and is distributed worldwide. It attacks more than 100 plant species, including crops such as cotton and strawberry, vegetables, deciduous fruits, and ornamental plants^1^. *T. cinnabarinus* causes enormous economic damage due to reductions in both the yield and quality of affected crops, and is controlled by acaricide sprays^2,3,4^. Due to acaricide resistance and environmental pollution, there is an increasing demand for sustainable, environmentally friendly control methods. For example, phytoseiid mites have been used as effective biological control with of agents of spider mites on many cultivated crops^5, 6^. In this study, we tested the effectiveness of new bio-pesticides as alternatives to chemical pesticides for the control of strawberry mites. *Tetranychid* is an important pest that severely affects strawberry production in Pinggu district, Beijing. It can seriously damage leaves of strawberry plants, causing economic losses and potentially leading to death of the host plant.

Chemical pesticides are often applied to control strawberry mites. However, long-term use of one chemical can easily cause the mites to develop insecticide resistance, making them more difficult to control. Moreover, pick your own strawberry farms are an increasingly popular spring outing, so it is very important to produce strawberries free of artificial pesticides. Bio-pesticides are safe, organic, and tend not to induce resistance in target species.

Biological pesticides are the use of living beings and active ingredients of living beings, and natural compound of the material, the preparation of the prevention and control of plant diseases, insects and plant growth regulator. Compared with the traditional chemical pesticides, biological pesticides have some good features, as they are safe to living beings and non-target organisms, good environmental compatibility, not easy to produce resistance, easy to protect biodiversity, wide sources, etc. In recent years, many new biological resources have been used, get innovative production technology. Some new products come into the market^7^.

Relative study about the acaricidal mechanism was reported. Liang *et al*. ^8^ reported that the study on scopoletin toxicity to *Tetranychus cinnabarinus* and its acaricidal mechanism, several target enzymes activity of nervous system and the effect on acetylcholinesterase were tested. Wang *et al*. ^9^ reported that the colorimetry assay to test *Tetranychus cinnabarinus’* enzyme activity of few enzyme after applying the acaricidal activities of *Polygonum aviculare.* The purpose of our study was to explore the biological pesticides’ (veratrine, kanghebio) target enzyme of strawberry mites by enzymatic activity of Ca^2+^-adenosine triphosphatase, acetylcholinesterase and monoamine oxidase.

In this study, 13 acaricides were used, including newly listed chemical pesticides (spirotetramat,bifenazate), a new botanical preparation (veratrine), and microbial agents (kanghebio, spinosad). Laboratory toxicity tests and greenhouse experiments were carried out in order to evaluate the control efficacy of chemical pesticides and bio-pesticides, and explore the potential mechanisms by which they act.

## 2. MATERIALS AND METHODS

### 2.1. Selected Acaricides and test pest sources

In this experiment, 13 different acaricides were used and listed in Table 1. The experiment was carried out during Jan-May 2014.The tested mites were collected from strawberry greenhouses in Pinggu District, Beijing and kept in laboratories. Female adult mites were used for laboratory toxicity tests.

**Table 1.** 13 different acaricides were used in this experiment.

| Acaricides | Manufacturers |
| --- | --- |
| 10% Bifenthrinemulsifiable concentrate | Langfang Pesticides Medium-scale Trial Manufac<br>under IPP-CAAS |
| 1.8% Abamectinemulsifiable concentrate | Beijing C. A. H. Bio-technology Co., Ltd. |
| 15% Pyridabenemulsifiable concentrate | Jiangsu Kwin Co., Ltd. |
| 5% Hexythiazoxwetable powder | IPP pesticide factory of CAAS |
| 20% Carbosulfanemulsifiable concentrate | FMC Corporation |
| 2.5% Lambda-cyhalothrinemulsifiable concentrate | Syngenta Crop Protection Co., Ltd. |
| 24% Envidor suspending agent | Bayer Crop Science China Co., Ltd. |
| 2.5% Spinosad suspending agent | Dow Chemical Company |
| 20% Chlorantraniliprole suspending agent | USA Dupont Group Chemical Co., Ltd. |
| 10% Chlorfenapyr suspending agent | BASF SE |
| 0.5% Veratrine soluble liquid | Jianhua Plant Pesticide Plant,Handan City, Hebei |
| 43% Bifenazate suspending agent | Chemtura Inc. |
| Kanghebio | Korea Bio Co., Ltd. |

### 2.2. Test method

#### 2.2.1. Laboratory toxicity test

A slide-dip method^10,11^ based on the United Nations Food and Agriculture Organization (UN FAO) recommended method was used in laboratory toxicological experiments. The 3-cm lengths of double-sided adhesive tape were attached to the slides. Female mites, each with the same size, bright color, and high activity, were chosen, and the back of each mite was attached to the adhesive tape; between 15 and 25 mites were attached to each slide. There was a gradient of 5 concentrations of each pesticide. Each slide, along with attached mites, was dipped into a pesticide, shaken, removed from the pesticide after 5s, and then dried with filter paper. Then the slides were put in culture dishes with gauze, and these put in an incubator with the conditions used for acclimatizing the mites. The test was repeated three or four times for each pesticide, and slides with mites were dipped in water as a control. The number of live mites was counted after 6h, 12h, 24h, 36h, 48h, 60h, and 72h. Mites were checked with a small brush under a microscope; mites which did not move were considered dead.

#### 2.2.2. Greenhouse test

The greenhouse test was carried out in Pinggu district, Beijing. Two greenhouses (No.1 and No.2) were selected for the experiment; the variety of strawberry used was Sweet Charlie. Eight acaricides (abamectin, pyridaben, cyhalothrin, veratrine, carbosulfan, hexythiazox, bifenazate, and kanghebio) were tested, and all treatments were repeated three times. Twenty-eight replicated plots were set up. A back-mounted hand sprayer was used to spraying the acaricides. Ten strawberry plants were used in each experiment, and the number of strawberry mites on 5 leaves of each plant was counted before spraying. The number of remaining live strawberry mites was counted after 3d, 7d, 14d, and 21d using a microscope.

#### 2.2.3. Enzyme activity test

##### 2.2.3.1. Preparation of enzyme solution

Twenty-four hours after acaricide treatment, 100 female adult mites were homogenized with 0.25mL normal saline solution (NS) for 15min in a 4°C ice-water bath at 2500rpm,and the resulting supernatant was collected.

##### 2.2.3.2. Enzyme source protein concentration

G-250 Coomassie Brilliant Blue (5 mL) was mixed with enzyme liquid (0.2 mL) at 25°C. Optical density (OD) was measured three times at 595nm, and the protein content was calculated according to a standard curve.

#### 2.2.4. Acetylcholinesterase (TChE) test

The TChE checker board method was followed the instructions of enzyme activity test kit produced by Nanjing Jiancheng Bioengineering Institute. OD was measured three times at 412nm and zero set with double distilled water (DDW).

#### 2.2.5. Ca^2+^-ATPase test

The ultra-micro ATPase checkerboard method was followed the instructions of enzyme activity test kit produced by Nanjing Jiancheng Bioengineering Institute. OD was measured three times at 412nm and zero set with distilled water.

#### 2.2.6. Monoamine oxidase (MAO) test

The monoamine oxidase checkerboard method was followed the instructions of enzyme activity test kit produced by Nanjing Jiancheng Bioengineering Institute. OD was measured three times at 242nm and the enzyme activity calculated.

### 2.3. Data analysis

Comparisons of rates of pest mortality and revised control efficiency were conducted using one-way analysis of variance (one-way ANOVA). Significant differences among means were determined using Fisher’s LSD test at *P* = 0.05^12^, and all analyses were conducted in the statistical program SPSS v 19.0.

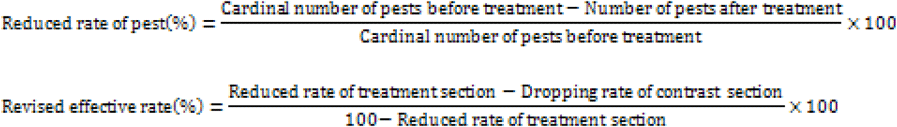

## 3. RESULTS

### 3.1. Laboratory test

The acaricide most effective in control of strawberry red spider mites was 1.8% abamectin, LC_50_ was 0.18 mg L^-1^ (Table 2). Following this, a group of five (15% pyridaben, 2.5% lambda-cyhalothrin, 0.5% veratridine, 20% carbosulfan) had high toxicity (LC_50_= 2.69 mg L^-1^, 3.94 mg L^-1^, 5.98 mg L^-1^ and 6.75 mg L^-1^, respectively). The next most toxic group was 5% hexythiazox and 10% bifenazate (LC_50_ = 9.82 mg L^-1^ and 19.09 mg L^-1^, respectively). The next group, 24% spirodiclofen, 20% chlorantraniliprole and 10% chlorfenapyr, was less effective; the order of toxicity from high to low was spirodiclofen>chlorantraniliprole>chlorfenapyr. The least effective group was 2.5% spinosadand 43% bifenazatewith LC50= 117.73mg L^-1^ and 146.22mg L^-1^, respectively.

**Table 2.** Toxicity of 13 acaricides on adult female strawberry mites using the slide-dip method in laboratory tests.

| Acaricide | Slope ± SE | LC<sub>50</sub>(ng a.i.adult<sup>-1</sup>) | 95% confidence interval<br>(ng a.i.adult<sup>-1</sup>) | $\chi^2$(df) | Relative toxicity index |
| --- | --- | --- | --- | --- | --- |
| Abamectin | 1.29±0.15 | 0.18 | 0.14–0.25 | 1.23(16) | 0.0012 |
| Pyridaben | 0.79±0.10 | 2.69 | 1.55–4.48 | 2.35(19) | 0.0184 |
| Cyhalothrin | 1.85±0.35 | 3.94 | 3.01–5.11 | 5.68(16) | 0.0269 |
| Veratrine | 1.68±0.20 | 5.98 | 4.83–7.53 | 12.57(13) | 0.0409 |
| Carbosulfan | 1.53±0.19 | 6.75 | 5.24–8.85 | 1.90(16) | 0.0462 |
| Hexythiazox | 2.18±0.28 | 9.82 | 7.89–11.79 | 5.48(19) | 0.0672 |
| Bifenthrin | 1.17±0.25 | 19.09 | 11.79–27.49 | 2.33(16) | 0.1306 |
| Envidor | 1.84±0.34 | 40.19 | 30.98–52.49 | 4.44(16) | 0.2749 |
| Chlorantraniliprole | 0.96±0.17 | 60.50 | 38.74–92.85 | 3.45(16) | 0.4138 |
| Chlorfenapyr | 2.89±0.54 | 75.84 | 65.31–91.40 | 4.22(16) | 0.5187 |
| Spinosad | 0.92±0.16 | 117.73 | 77.47–197.82 | 2.09(16) | 0.8052 |
| Bifenazate | 1.97±0.21 | 146.22 | 120.82–180.02 | 4.98(13) | 1.00 |
| Kanghebio | 1.18±0.19 | 4.15$\times 10^7$ | 2.54$\times 10^7$–<br>5.73$\times 10^7$ | 1.12(13) | |

We tested 13 acaricides, including chemical agents (e.g., abamectin, bifenazate) a botanical pesticide (veratrine) and a microbial agent (kanghebio). The most effective was abamectin. Three new acaricides (veratrine, kanghebio and bifenazate) had variable effectiveness. Veratrine and kanghebio were more effective than bifenazate. Sixacaricides with high efficacy were identified: abamectin, pyridaben, cyhalothrin, carbosulfan, hexythiazox, and bifenthrin.

The status of female mites 24h after acaricide was determined under a fluorescence microscope. The status of female mites treated with avermectin, pyridaben, kanghebio and cyhalothrin changed significantly; they curled up in convulsions, and their activity was low. Female mites treated with kanghebio became black. Female mites treated with veratridine, carbosulfan, hexythiazox and bifenthrin, curled their feet and had low activity, but no convulsions. After treatment with chlorfenapyr, mites were black, with curled feet and no convulsions or significant change in the activity. There were no significant differences among female mites treated with acaricide (spirotetramat, chlorantraniliprole, spinosad, and bifenazate) and those treated with water as a control. Figure 1 showed the proportional reduction in numbers of live mites 6 h to 72 h after the slide-dip test. Abamectin and pyridaben had the most rapid reduction in mite numbers (i.e.,6h after the test),while spinosad, chlorfenapyr, chlorantraniliprole and bifenazate had little effect after 6h. Pyridaben and carbosulfan showed the greatest reductions in mite numbers at 24h and 48h after the test. Bio-pesticides (veratrine, spinosad, and kanghebio) performed worse than chemical pesticides. Most acaricides’ effectiveness grew noticeably12h after the test, and all of them killed some mites between 1d and 3d.

**Figure 1.**
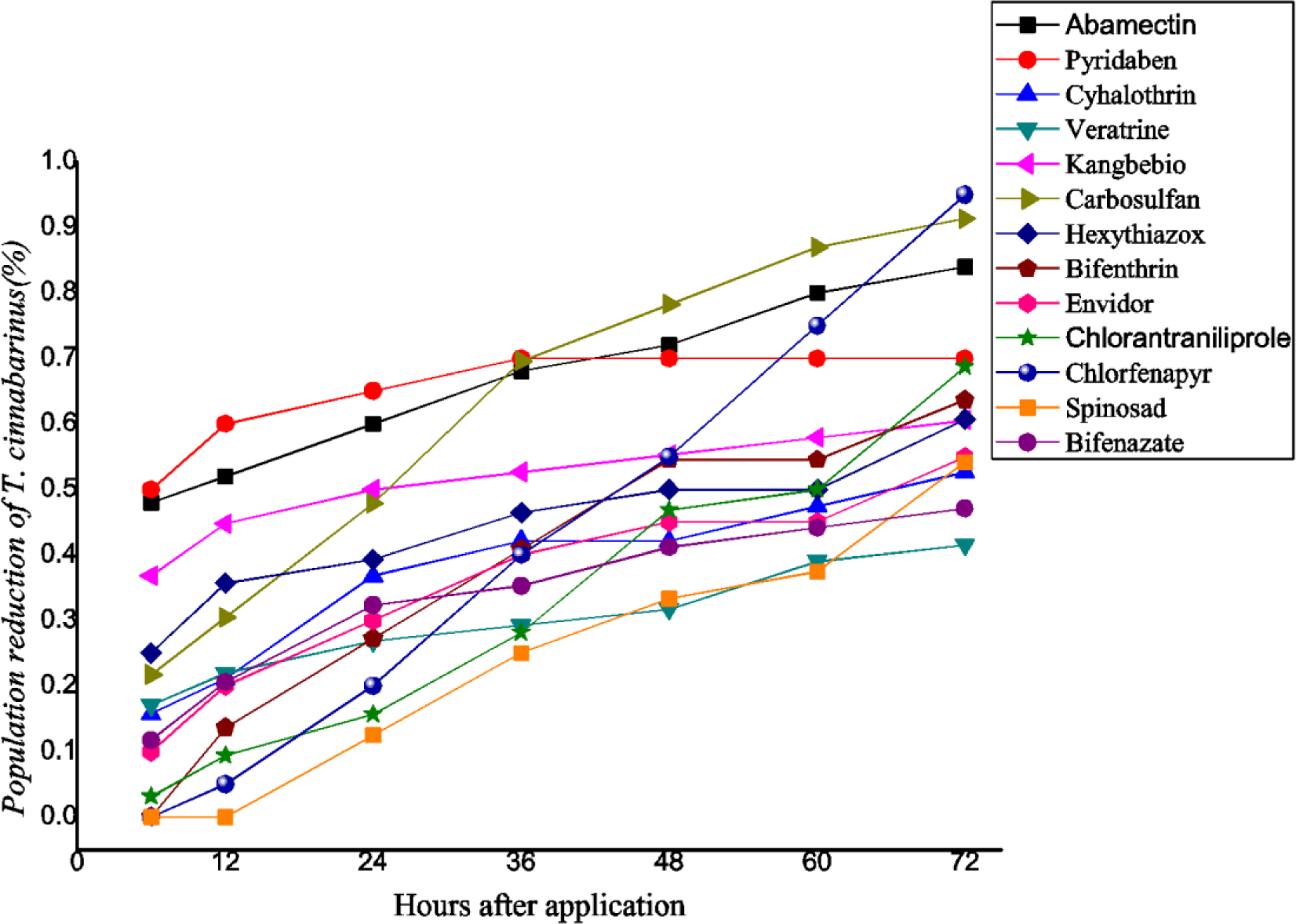
Reductions in numbers of adult female mites following treatment with one of 13 acaricides in slide-dip laboratory tests (6–96h).

Figure 2 showed the status of female mites 24 h after acaricide application under a fluorescence microscope. The status of female mites treated with abamectin, pyridaben, kanghebio and cyhalothrin (panels A, B, C and D, respectively)was greatly changed-many were curled up in convulsions and activity was low. Female mites treated with kanghebio (C) became black. Panels E, F, G, and H show female mites treated with veratrine, carbosulfan, hexythiazox, and bifenthrin, respectively. Mites had curled feet and low activity but no convulsions. Panel K (treatment with chlorfenapyr): mite black, foot curled, no convulsions, no significant change in activity. Compared to the control (water, Panel CK), female mites treated with spirotetramat, chlorantraniliprole, spinosad and bifenazate (Panels I, J, L, and M, respectively) showed no significant change in status.

**Figure 2.**
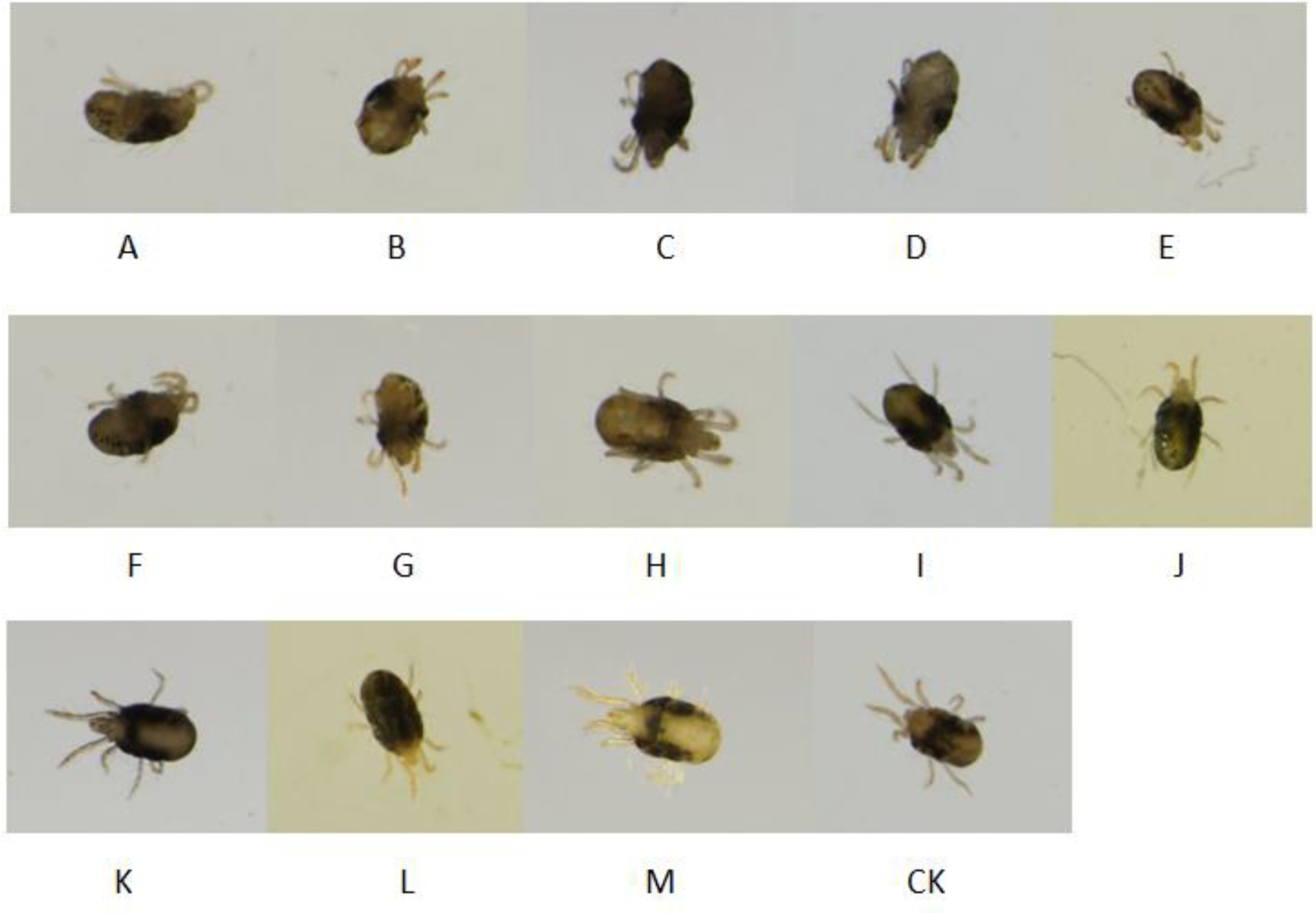
The status of female adult mites 24 h after acaricide application, viewed under a stereo fluorescence microscope. Note: Treated with A(Abamectin), B(Pyridaben), C(Kanghebio), D(Cyhalothrin), E(Veratrine), F (Carbosulfan), G(Hexythiazox), H(Bifenthrin), I(Envidor), J(Chlorantraniliprole), K(Chlorfenapyr), L(Spinosad), M(Bifenazate), CK(Water).

### 3.2. Greenhouse test

The same treatments were applied in No.1 and No. 2 greenhouses over the same period. The overall trend was consistent with the laboratory test (Table 3 and 4). Two new bio-pesticides showed better control efficacy than chemical pesticides in greenhouse tests. Of the eight acaricides, abamectin showed the best efficacy, while Pyridaben, kanghebio, cyhalothrin, veratrine, carbosulfan, hexythiazox, and bifenthrin also showed good efficacy. The results of experiments in No.1greenhouse were consistent with those from No.2 greenhouse, and these results were in agreement with those of the laboratory test. However, better control was achieved in No.2 greenhouse than in No.1greenhouse. This was possibly due to moister conditions in No.2 greenhouse, which were not suitable for red spider outbreak, while conditions were more suitable for red spider in the No. 1 greenhouse. Eight acaricides were effective in prevention and control, and the efficacy of the biological agents veratrine and kanghebio was good. Combined with laboratory results, this suggests that the biological acaricides veratrine and kanghebio, which are low in toxicity and high effective, could be applied greenhouses to control red spider.

**Table 3.** Reductions in insect density and corrected mortality results for 8 acaricides from the experiment in Greenhouse 1.

| Acaricide | Reduction in insect density (%) |  |  |  | Correctedmortality (%) |  |  |  |
| --- | --- | --- | --- | --- | --- | --- | --- | --- |
|  | 7d | 14d | 21d | 28d | 7d | 14d | 21d | 28d |
| 1.8% Abamectin EC | 91.93 | 93.57 | 95.75 | 97.14 | 98.28ab | 98.77ab | 99.34a | 99.68a |
| 15% Pyridaben EC | 92.63 | 94.75 | 95.63 | 96.03 | 98.44a | 99.08a | 99.31a | 99.50a |
| Kanghebio | 84.30 | 88.08 | 90.46 | 91.45 | 96.87bc | 98.01abc | 98.59ab | 98.97ab |
| 2.5% Cyhalothrin EC | 81.29 | 83.03 | 84.20 | 88.63 | 95.92c | 96.65bc | 97.41b | 98.50bc |
| 0.5% Veratrine SL | 80.91 | 81.58 | 83.64 | 84.19 | 95.85c | 96.41bc | 97.31b | 97.91bc |
| 20% Carbosulfan EC | 77.06 | 78.48 | 80.38 | 81.04 | 95.02c | 95.78c | 96.88b | 97.57c |
| 5% Hexythiazox WP | 76.33 | 77.87 | 78.57 | 79.28 | 94.91c | 96.21bc | 96.68b | 97.30c |
| 10% Bifenthrin EC | 49.63 | 32.46 | 15.55 | 11.13 | 89.02d | 84.37d | 86.45c | 88.48d |

**Table 4.** Reductions in insect density and corrected mortality results for 7 acaricides from the experiment in Greenhouse 2.

| Acaricide | Reduction in insect density % |  |  |  | Correctedmortality % |  |  |  |
| --- | --- | --- | --- | --- | --- | --- | --- | --- |
|  | 7d | 14d | 21d | 28d | 7d | 14d | 21d | 28d |
| 1.8% Abamectin EC | 94.03 | 95.03 | 96.80 | 98.14 | 97.90a | 98.75a | 99.42a | 99.86a |
| 15% Pyridaben EC | 93.54 | 95.62 | 96.23 | 97.27 | 97.73a | 98.90a | 99.32a | 99.61ab |
| Kanghebio | 86.20 | 88.81 | 90.46 | 91.80 | 95.36b | 97.53ab | 98.37ab | 98.90bc |
| 2.5% Cyhalothrin EC | 82.59 | 84.18 | 86.09 | 90.18 | 93.73bc | 95.94bc | 97.37bc | 98.52cd |
| 0.5% Veratrine SL | 82.56 | 83.57 | 85.39 | 86.04 | 93.76bc | 95.85bc | 97.22bc | 97.89cde |
| 20% Carbosulfan EC | 77.40 | 79.59 | 80.60 | 81.52 | 91.89c | 94.77bc | 96.44bc | 97.29de |
| 5% Hexythiazox WP | 75.46 | 76.75 | 76.75 | 77.03 | 91.23c | 94.28c | 95.80c | 96.60e |
| 10% Bifenthrin EC | 50.45 | 35.10 | 17.33 | 5.85 | 82.17d | 80.22d | 84.59d | 86.10f |

### 3.3. Determination of enzyme activity

Enzyme activity tests were carried out to explore possible mechanisms of action. Kanghebio and cyhalothrin obviously inhibited Ca^2+^-adenosine triphosphatase (Ca^2+^-ATPase), while veratrine and kanghebi oobviously inhibited acetylcholinesterase (TChE) and monoamine oxidase (MAO). Cyhalothrin exerted an auxo-action on MAO.

Enzyme activity results were shown in table 5. Three acaricides showed different effects on TChE, Ca^2+^-ATPase and MAO. Kanghebio and cyhalothrin obviously inhibited Ca^2+^-ATPase, while the two Bio-pesticides (veratrine and kanghebio) obviously inhibited TChE and MAO. Cyhalothrin showed auxo-action on MAO.

**Table 5.** Enzyme activity of female adult mites 24h after treatment with acaricides by acetylcholinesterase checkerboard, ultramicroatpase checkerboard and monoamine oxidase checkerboard.

| Acaricide | Ca<sup>2+</sup>-ATPase(*Umgprot<sup>-1</sup>*) |  | TChE(*Umgprot<sup>-1</sup>*) |  | MAO(*Umgprot<sup>-1</sup>*) |  |
| --- | --- | --- | --- | --- | --- | --- |
|  | Dose(*mg a.i. L<sup>-1</sup>*) |  | Dose(*mg a.i. L<sup>-1</sup>*) |  | Dose(*mg a.i. L<sup>-1</sup>*) |  |
|  | 2.5 | 0.625 | 500 | 125 | 100 | 25 |
| 0.5% Veratrine EC | 4.9481 | 5.5596 | 0.0204 | 0.0990 | 0.0000 | 0.0034 |
| Kanghebio | 0.9043 | 1.5281 | 0.0205 | 0.0997 | 0.0091 | 0.0101 |
| 2.5% Cyhalothrin EC | 1.4034 | 3.1075 | 0.0353 | 0.1457 | 0.0341 | 0.0265 |
| CK(DDW) | 6.5490 |  | 0.1605 |  | 0.0114 |  |

Figure 3 shows the inhibiting effects of three acaricides on AChE. The bio-pesticides (veratrine, kanghebio) had the same inhibiting effect on AChE, veratrine and kanghebio’s inhibiting effects were greatest at high concentrations and cyhalothrin’s inhibiting effect was lowest at low concentrations.

**Figure 3.**
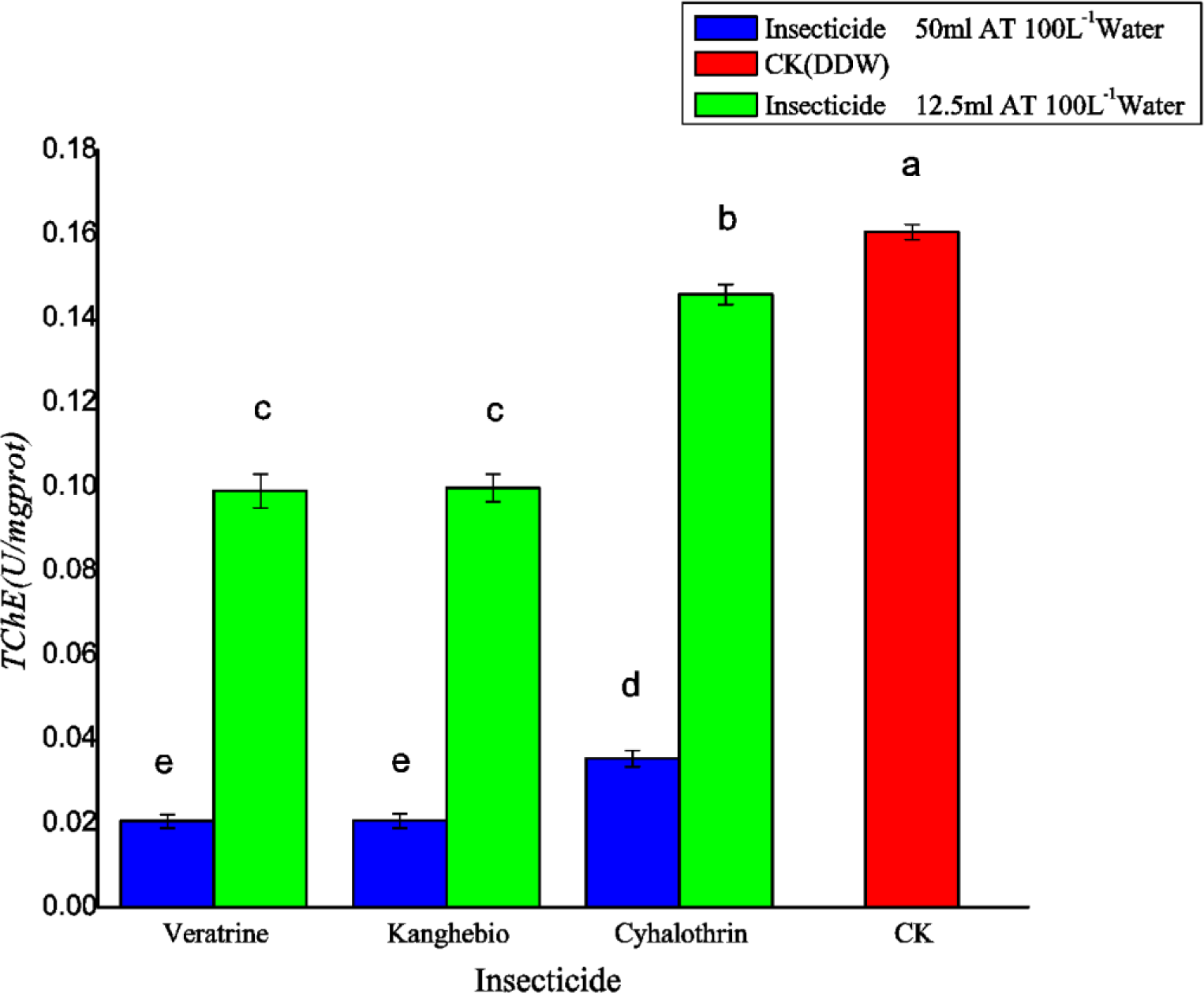
Acetylcholinesterase activity of female adult mites 24h after treatment with one of three acaricides.

Figure 4 shows the inhibiting effects of three acaricides on ATPase. Kanghebio had the greatest inhibiting effect, followed by cyhalothrin, while veratrine had the least inhibiting effect.

**Figure 4.**
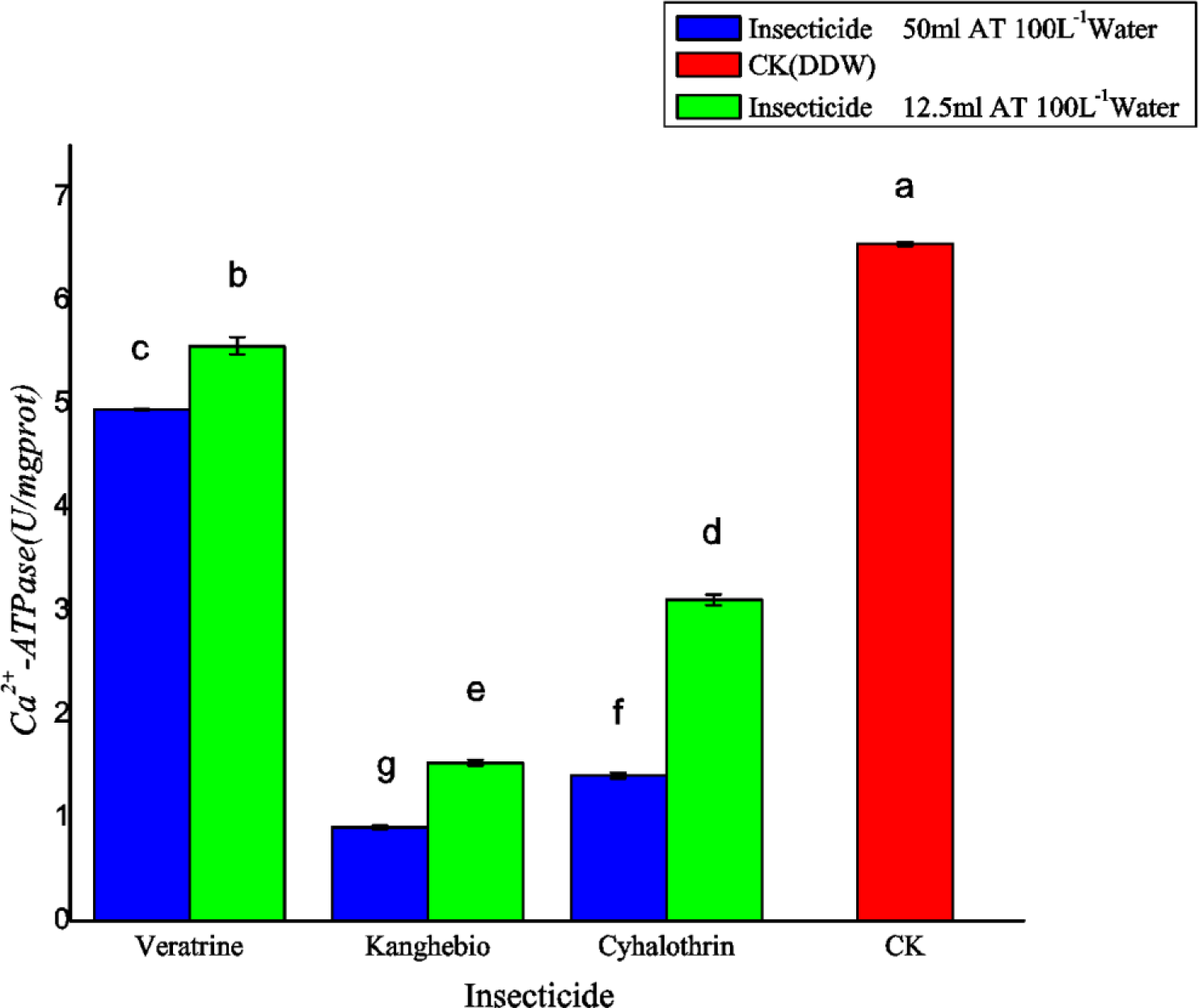
Ca2+-ATPaseactivity of female adult mites 24h after treatment with one of three acaricides.

Figure 5 shows the inhibiting effect on MAO of two acaricides (veratrine and kanghebio). At high concentrations veratrine had an obviously inhibiting effect, enzyme activity approached zero. Cyhalothrin hadauxo-action on MAO, and this was greatest at high concentrations of cyhalothrin.

**Figure 5.**
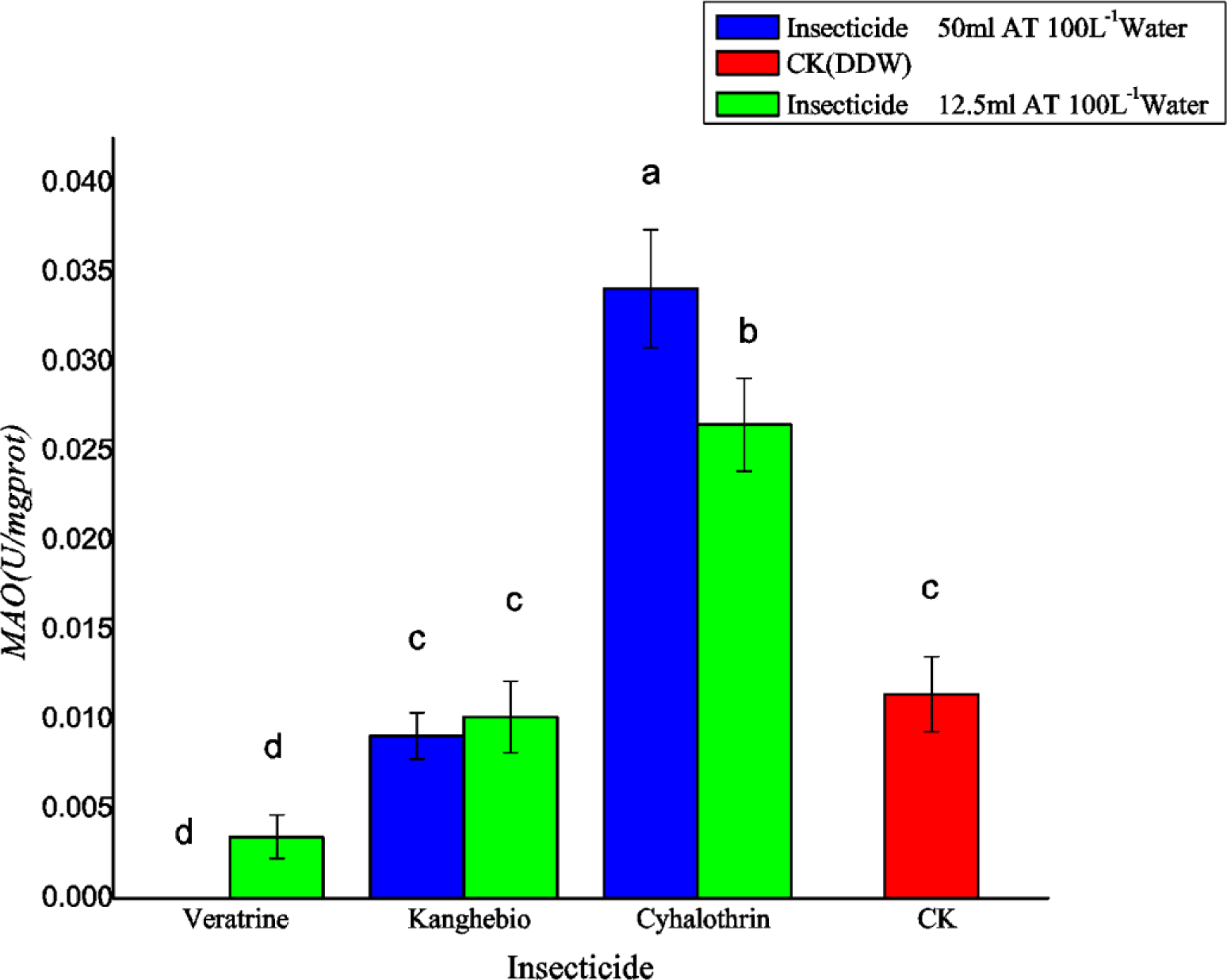
Monoamine oxidase activity offemale adult mites 24h after treatment with one of three acaricides.

## 4. DISCUSSION

Strawberry mite infestations often occur during fruiting, mites can rapidly develop resistance to numerous acaricides, this adaptation can give rise to rapid population increases^13,14^. The dominant species of spider mite is *T. cinnabarinus*, which can greatly influence the quality and yield of strawberry crops.

The results showed as follows: of the 13 acaricides tested, those with better efficacy in the control of strawberry mites were Abamectin, Pyridaben, cyhalothrin, veratrine, and kanghebio. Veratrine and kanghebio are bio-pesticides and were as effective as conventional acaricides. They are environment-friendly and effective acaricides with low toxicity. Next, 0.5% of the recommended doses of veratrine and kanghebio are 100mlha^-1^ and 150mlha^-1^, respectively. It is recommend that pesticide be applied in early stages of insect attack, spraying once every 3d during severe infestations and spraying once every 7d after insect numbers decrease, then spraying once every 10d for prevention.

Spider mites were sensitive to abamectin and have not become resistant to abamectin in the Pinggu area. Some reports^15-19^suggest that strawberry mites have developed pesticide resistance. Therefore, we do not recommend abamectin as a long-term option for controlling mites. Of the chemical acaricides, pyridabenand cyhalothrin showed high efficacy and may have unwanted effects on bees during strawberry flowering. They are WHO Class II and not environmentally friendly, so we do not recommend that they be widely applied in the field.

Carbosulfan and hexythiazox have relatively lower toxicity and were ineffective at low concentrations. This finding suggests the dosage should be increased to improve the efficacy. Such acaricides that do not meet the requirements of low toxicity and high efficacy are not useful for preventing strawberry spider mites in greenhouses.

Combining the results of the laboratory test with those of the greenhouse experiment, we recommend two bio-pesticides (veratrine and kanghebio) and one chemical pesticide (cyhalothrin) to control female adult mites, explore the possible role targets by acetylcholinesterase checkerboard, ultramicroatpase checkerboard and monoamine oxidase checkerboard. The results showed that the two bio-pesticides (veratrine and kanghebio) inhibited Ca^2+^-ATPase, TChE and MAO. The chemical pesticide (cyhalothrin) inhibited Ca^2+^-ATPase, TChE and had auxo-action on MAO. We speculate that Ca^2+^-ATPase and TChE inhibition is the mechanism by which these acaricides act.

The results obtained in our study show that the bio-pesticides veratrine, kanghebio are effective control agents both in laboratory tests and greenhouse experiments. Furthermore, they are environmentally friendly acaricides, with low toxicity and high efficacy. This research provides a strong basis for using bio-pesticides in place of conventional chemicals to control strawberry spider mites, and provides useful insights into the biological mechanisms by which acaricides act. However, we need to further study their efficacy at different mite developmental stages and clarify the mechanisms of action of these bio-pesticides.

## 5. CONCLUSION

In summary, two new bio-pesticides and several chemical pesticides were compared using laboratory toxicity tests and greenhouse experiments. The bio-pesticides showed better control efficacy than chemical acaricides in greenhouse tests. These bio-pesticides have developed a good reputation in international markets in recent years due to their safety, organic nature, environmental friendliness, and lack of propensity to induce pesticide resistance, among other advantages. Our study provides a scientific basis for chemical pesticides to be replaced by these and potentially other new bio-pesticides.

## ACKNOWLEDGMENTS

This research was supported by National Natural Science Foundation of China (Program No. 31672066), Beijing Innovation Consortium of Agriculture Research System(BAIC01-2017), and National Key Research and Development Program of China(2018YFD0201300).

## REFERENCES

[1] L.R. Jeppson, H.H. Keifer, E.W. Baker, Mites injurious to economic plants. Univ of California Press, (1975).

[2] T.F. Akşit, Özsemerci, İ Çakmak, Studies on determination of harmful fauna in the fig orchards in Aydın province (Turkey). TurkEntomol Derg 27181–189(2003).

[3] İ Çakmak, H. Başpınar, Studies on insect pests and mites and their natural enemies on summer vegetables in Aydın province. Aegean Region 1st Agricultural Congr; Aydın, Turkey, pp. 427–435(1998).

[4] İ Çakmak, H. Başpınar, N. Madanlar, The population densities of spider mites and their natural enemies on protected strawberries in Aydın province. Turk Entomol Derg 27:91–205 (2003).

[5] W. Helle, M.W. Sabelis, Spider Mites: Their Biology, Natural Enemies and Control. Vol. 1B. Elsevier, Amsterdam, the Netherlands, (1985).

[6] J.C. VanLenteren, M. Benuzzi, G. Nicoli, S. Maini, Biological control in protected crops in Europe.In: Biological Control and Integrated Crop Protection: Towards Environmentally Safer Agriculture (Eds. van Lenteren J.C., A.K. Minks and O.M.B. de Ponti), Pudoc, Wageningen, The Netherlands, pp. 77–89(1992).

[7] B.G. Gu, H. Jiang, The industrialization of microbial pesticides. Science Press, 13–18(2000).

[8] W. Liang, X.N. Bai, L.Q. Ma, G.L. Shi, Y.N. Wang, Preliminary study on scopoletin toxicity to Tetranychuscinnabarinus and its acaricidal mechanism. ScientiaSilvae Sinicae 8:68–71(2011).

[9] Y.N. Wang, Q. Li, Z.H. Lu, G.L. Shi, Acaricidal Activities of Polygonumaviculare Extracts against Tetranychus cinnabarinus and Their Effects on Enzyme Activities in the Mite. Scientia Silvae Sinicae, 46:3103–107 (2010).

[10] J.R. Busvine, Recommended methods for measurement of resistance to pesticides. FAO Plant Production and Protection Paper, 21:349–54(1980).

[11] Y.Q. Zhang, W. Ding, J. Wu, Z. Zhao, Comparison of three acaricide bioassay methods. Pesticide Science and Administration, 28:350–53(2007).

[12] R.G.D. Steel, J.H. Torrie, Principles and procedures of statistics. America: McGraw-Hill Book Co., Inc. New York, p 481 (1960).

[13] D. Bernardi, M. Botton, U.S. Cunha, O. Bernardi, T. Malausa, M.S. Garcia, D.E. Nava, Effects of azadirachtin on Tetranychusurticae (Acari: Tetranychidae) and its compatibility with predatory mites (Acari: Phytoseiidae) on strawberry. Pest management science, 69:375–80(2013).

[14] I. Çakmak, Başpinar N. Madanlar, Control of the carmine spider mite Tetranychus cinnabarinus Boisduval by the predatory mite Phytoseiuluspersimilis (Athias-Henriot) in protected strawberries in Aydin, Turkey, Turkish journal of agriculture and forestry, 29:3259–265(2005).

[15] Y.J. Gong, Z.H. Wang, B.C. Shi, W.X. Gui, G.H. Jin, Y.Y. Sun, S.J. Wei, Sensitivity of different field population of Tetranychusurticae Koch (Acari: Tetranychidae) to the acaricides in beijing area. Scientia Agricultura Sinica, 47:32990–2997 (2014).

[16] M. Castagnoli, M. Liguori, S. Simoni, C. Duso, Toxicity of some insecticides to Tetranychus urticae, Neoseiulus californicus, and Tydeus californicus. Bio Control, 50:3611–622(2005).

[17] C. Kazak, K. Karut, I. Kasap, C. Kibritci, E. Sekeroglu, The potential of the hatay population of Phytoseiuluspersimilis to control the carmine spider mite Tetranychuscinnabarinus in strawberry in Silifke-Icel, Turkey. Phytoparasitica, 30:3451–458 (2002).

[18] C. Kazak, C. Kibritci, Population parameters of Tetranychus cinnabarinus Boisduval (Prostigmata: Tetranychidae) on eight strawberry cultivars. Turkish Journal of Agriculture and Forestry, 32:319–27(2008).

[19] Y. J. Luo, J. Ni, Y.G. Liu, X.J. Jiang, J.P. Chai, D.Y. Xie, A.S. Da, P. Huang, X. L. Shao, W. Ding, Acaricide resistance of Tetranychus cinnabarinus (Acari: Tetraychidae) from mulberry plantations in southwest China. International Journal of Acarology, 39:3522–525(2013).

